# MotifAE Reveals Functional Sequence Patterns from Protein Language Model: Unsupervised Discovery and Interpretability Analysis

**DOI:** 10.1101/2025.11.04.686576

**Authors:** Chao Hou, Di Liu, Yufeng Shen

## Abstract

Protein language models (pLMs) learn sequence patterns at evolutionary scale, but these patterns remain inaccessible within these “black box” models. To discover them, we developed MotifAE, an unsupervised framework based on the sparse autoencoder (SAE) architecture that projects pLM embeddings into an interpretable, sparse latent space. MotifAE introduces an additional smoothness loss to encourage coherent feature activation, which markedly improves the identification of known functional motifs compared to the standard SAE. The sequence patterns captured by MotifAE exhibit rich diversity, align with known functional motifs, and are reflected in the model’s weight space. Beyond short motifs, MotifAE also captures structural domains, with latent feature activation scores correlating with residue importance for different domain functions. By aligning MotifAE features with experimental data, we further identified features associated with domain folding stability. These features enable the prediction of a stability-specific fitness landscape that improves stability prediction and supports the engineering of domains with enhanced stability. Overall, MotifAE provides a general framework for systematic sequence pattern discovery and interpretation, with the potential to advance protein function analysis, mutation effect interpretation, and rational protein engineering.

## Introduction

Proteins mediate essential biological functions. Their functional elements, such as catalytic centers, protein/nucleic acid/small molecule binding interfaces, and post-translational modification sites, exhibit conserved sequence patterns commonly referred to as motifs. Experimental identification of a motif requires first locating protein regions that share the same function and then aligning them to get the sequence pattern, which is both labor-intensive and time-consuming. As of 2025, only ∼350 motifs have been curated in the Eukaryotic Linear Motif (ELM) database^1^. To complement experimental efforts, machine learning methods have been developed^2–4^, often trained on curated resources like ELM. However, these supervised models are typically tailored to specific functions and inherently biased toward their training datasets, limiting their generalizability. Therefore, systematic and unbiased discovery of functional motifs is crucial for advancing our understanding of protein functions.

Besides short motifs, structural domains also exhibit conserved sequence patterns. By aligning experimentally solved structures in the Protein Data Bank^5^ (PDB) and high-quality predicted structures, domain sequence patterns have been systematically characterized^6,7^. However, the functional organization within domains remains poorly understood. In particular, it is unclear how individual residues contribute to folding stability, catalytic activity, partner binding, or other functional roles. Clarifying residue-level importance across these functions is essential for understanding domain organization and has the potential to enable the rational engineering of domains with desired properties.

In recent years, deep learning has driven remarkable advances in modeling protein sequence^8^, structure^9^, and function^10^. A major development is protein language models (pLMs), such as ESM2^8^, which are self-supervised models trained on large-scale protein databases using masked or next-token prediction. Through this pre-training, pLMs implicitly learn conserved sequence patterns at evolutionary scale. For example, studies have shown that pLM predicts protein structure by capturing pairwise contact motifs^11^, pLM attentions identify structural contacts and target binding sites^12^, and pLM embeddings show similarity for proteins within the same domain family^13^. Moreover, numerous studies have leveraged pLM embeddings to predict functional regions such as protein sorting signals, transmembrane regions and post-translational modification sites^3^. Collectively, these studies demonstrate that pLMs encode rich information about the sequence patterns of functional motifs and structural domains. However, because pLMs operate as “black boxes”, the sequence patterns they encode is not directly accessible.

Various attempts have been made to extract interpretable features from language models. Among them, sparse autoencoders (SAEs) have shown strong potential for interpretability^14^. The core assumption of SAEs is that the model embedding can be expressed as a linear combination of distinct features^15^, with each feature ideally corresponding to an interpretable underlying factor. To achieve this, SAE uses an encoder to project embeddings into a higher-dimensional, sparse latent space, and uses a decoder to reconstruct the embeddings from this space. During SAE training, a reconstruction loss was used to ensure that the embeddings are accurately reconstructed, and a sparsity constraint was used to encourage sparse feature activations. Hereafter, we refer to each neuron in the SAE latent space as a feature, and neurons with positive values are activated. Typically, only tens of features are active per residue, far fewer than the original embedding dimension, which aids interpretability. SAEs have been successfully applied in large language models (LLMs) to find interpretable semantic features^14,15^. Adapting the training and interpretation strategies developed for LLM SAE, recent works found that applying SAE to pLMs has the potential to identify functional regions and structural elements^16–18^. However, it remains unclear whether biologically informed priors can improve pLM SAE interpretability and whether SAEs can systematically reveal motif sequence patterns and domain functional organization.

Here, we developed MotifAE, introducing methodological advances in both SAE training and interpretation to better suit it for biological sequences and compatibility with experimental data. We incorporated an additional smoothness loss to encourage MotifAE’s latent features to activate coherently, which markedly improves the identification of known functional motifs compared to the standard SAE. To analyze the motif sequence patterns captured, we computed Position-Specific Scoring Matrices (PSSMs) for MotifAE features, which reveals rich sequence diversity and highlights MotifAE’s potential for systematic functional motif discovery. For longer sequence patterns, MotifAE significantly outperforms the standard SAE in domain identification, with latent feature activation scores correlating with residue importance for different domain functions. Furthermore, by supervised alignment with experimental data, we identified MotifAE features associated with domain folding stability; these features subsequently improve performance in domain stability prediction and enable rational protein engineering with enhanced stability, as evaluated *in silico*. Overall, MotifAE provides a systematic framework for functional sequence pattern discovery from pLMs and annotation using curated databases and experimental data.

## Results

### MotifAE is a sparse autoencoder with coherent latent feature activation

We developed MotifAE, an adaptation of the sparse autoencoder (SAE) for discovering functional sequence patterns from pLMs. A standard SAE is trained with a reconstruction loss and a sparsity constraint (**Figure 1**), ensuring that the original embeddings are accurately reconstructed while enforcing sparse activation of latent features. To enhance biological relevance, we introduced a smoothness loss (see Methods), which requires that at least one neighboring residue within a defined window exhibits latent feature activations similar to the current residue (**Figure 1**; see Methods), thereby promoting coherent feature activation along the sequence. The neighboring residues with similar activations are not necessarily directly adjacent. This design is motivated by the observations that protein motifs often involve contiguous residues, that basic structural elements exhibit characteristic local patterns, such as alternating interactions in β-strands and periodic interactions in α-helices, and that structural domains are assembled from functional motifs and basic structural elements. A window size of three residues was used in this study. Because the smoothness loss is computed over the entire sequence, it can propagate along the sequence to capture longer-range relationships, extending beyond the local window.

**Figure 1.**
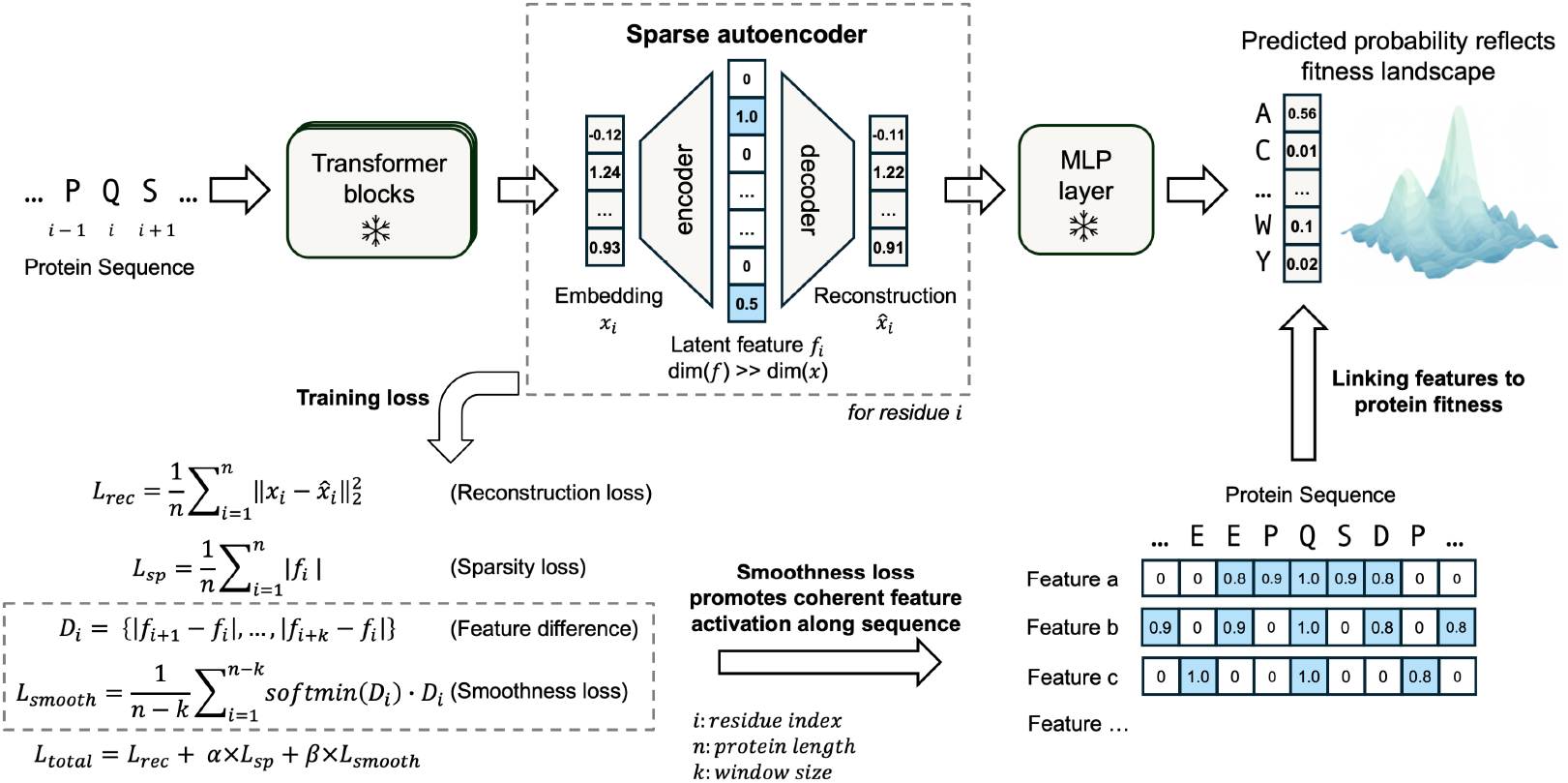
The MotifAE framework. MotifAE uses the sparse autoencoder (SAE) architecture with an additional smoothness loss. MotifAE projects ESM2 embeddings (dimension 1,280) into a high-dimensional latent space (dimension 40,960) using an encoder, and then reconstructs the embeddings through a decoder. The reconstructed embeddings can be projected to amino acid probabilities via the fixed ESM2 MLP layer, enabling prediction of the protein fitness landscape. MotifAE is trained with three objectives: a reconstruction loss, a sparsity loss, and a smoothness loss, the latter encourages coherent latent feature activations.

We trained MotifAE on the final-layer embeddings from ESM2-650M, which has shown strong performance across diverse downstream tasks. ESM2-650M captures the protein fitness landscape and is widely used to predict mutation effects^19–21^. We used the last-layer embeddings because they are directly used to predict amino acid probability which reflects the protein fitness landscape^19,20^, allowing us to investigate the relationship between latent features and fitness (**Figure 1**). To ensure that MotifAE captures representative protein features, training was performed on 2.3 million representative proteins obtained from structure-based clustering of the AlphaFold2 structure database^22^. The dimension of the latent space and weights for different losses were determined by jointly considering reconstruction error, latent sparsity, and the protein fitness prediction performance (**Figure S1**; see Methods). Based on these criteria, we selected a hidden dimension of 40,960, corresponding to a 32-fold expansion over the ESM2 embedding dimension. Notably, we found that the reconstruction of ESM2 embeddings is highly sensitive: although the reconstruction errors show little variation across different latent space dimensions (**Figure S1B**), the quality of the reconstructed embeddings varies substantially, as reflected in their performance on protein fitness prediction (**Figures S1D–E**).

We trained a standard SAE without the smoothness loss for comparison (see Methods). Both MotifAE and SAE achieve low reconstruction errors (**Figure 2A**), with approximately 46 latent features activated per residue on average in both models (**Figure 2B**). To assess the quality of the reconstructed embeddings, we evaluated their performance on protein fitness prediction. We projected the reconstructed embeddings through the fixed ESM2 MLP layer to predict amino acid probability (**Figure 1**), calculated the wild-type marginal log-likelihood ratio (LLR; see Methods)^19,21^, and used the LLR to estimate mutation effects. Across deep mutational scanning (DMS) experiments in ProteinGYM^20^ and pathogenic/benign mutations in ClinVar^23^ (see Methods), reconstructions from both MotifAE and SAE perform comparably to the original ESM2 embeddings (**Figure 2C-D**). The smoothness loss of MotifAE was introduced to encourage coherent latent feature activation. To evaluate this, we first analyzed the normalized difference in feature activation between neighboring residues and found that MotifAE features were more similar among nearby residues than those learned by the standard SAE (**Figure 2E**). We further analyzed the length distribution of activations and found that MotifAE features more frequently form clustered and contiguous activations than those of SAE (**Figure 2F**). Activated regions with a length of 5–30 residues suggest possible functional motifs, while those longer than 30 residues suggest possible domains.

**Figure 2.**
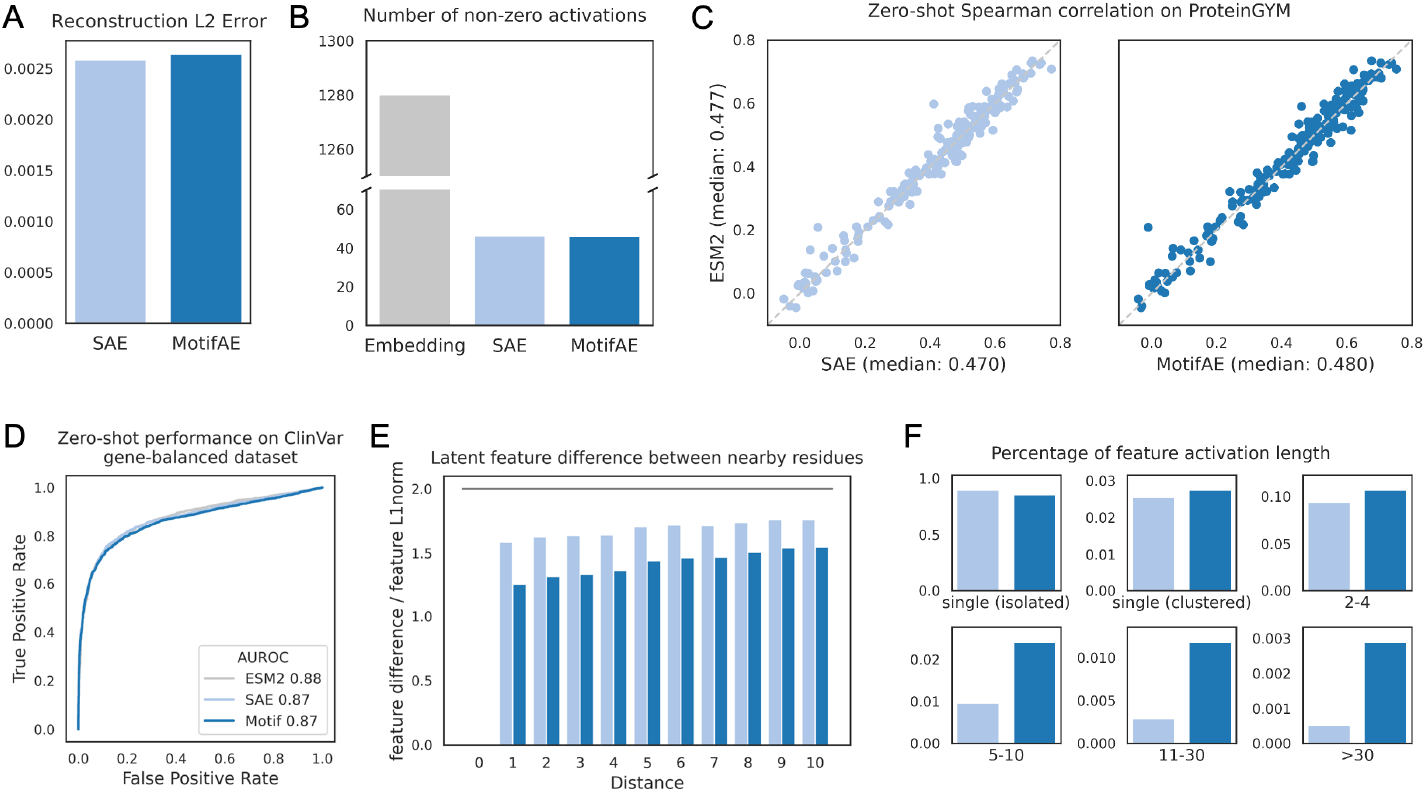
Comparison of MotifAE and SAE in terms of reconstruction error, latent sparsity, fitness prediction performance, and activation length distribution. (**A**) ESM2 embedding reconstruction error on the evaluation set (see Methods). (**B**) Mean number of non-zero neurons per residue. The ESM2 embedding has 1280 non-zero neurons, while both SAE and MotifAE have ∼46 non-zero (positive) latent features per residue. (**C**) Zero-shot performance (spearman correlation) on ProteinGYM DMS datasets, only single substitution mutations were evaluated. Each dot represents a DMS dataset. Wild-type marginal log-likelihood ratio (LLR) was used to estimate mutation effects. (**D**) Zero-shot performance on classifying pathogenic and benign missense mutations in the ClinVar gene-balanced dataset (see Methods). (**E**) Latent feature difference between nearby residues with different distances, measured using the absolute activation difference divided by the L1 norm of the activations. The value on y-axis is 0 for identical activations and ∼2 for random sparse activations. (**F**) Length distribution of activations from all features. For each feature, A single (clustered) activation is defined as an activated single residue with other activated residues within two amino acids along the sequence. In all plots, light blue denotes SAE, and dark blue denotes MotifAE.

Together, these results demonstrated that MotifAE preserves embedding reconstruction quality and latent space sparsity, while the smoothness loss promotes coherent feature activations, with the potential to enable the identification of functional motifs and structural domains.

### MotifAE captures known functional motifs

We first investigated whether MotifAE latent features capture functional motifs. Experimentally validated motifs were obtained from the ELM^1^ database, which covers six categories: ligand-binding, docking, post-translational modifications, targeting signals, degradation, and proteolytic cleavage sites. Each ELM motif contains multiple experimentally verified functional regions. For our analysis, we focused on 270 motifs with at least 20 residues in verified regions, as motifs with fewer annotated residues might be matched by chance due to the large number of latent features. We compared all MotifAE features against all ELM motifs, assessing whether the feature activation scores could distinguish residues located inside versus outside the experimentally verified motif regions (see Methods).

Considering the best-match feature for each motif, MotifAE demonstrates strong performance, achieving a median AUROC of 0.88 across the 270 motifs, significantly outperforming the standard SAE (median AUROC 0.80) (**Figure 3A**). The best-match MotifAE features achieve AUROCs above 0.8 for 193 motifs and above 0.9 for 114 motifs. One representative motif–feature pair is f13268 with the ELM motif MOD_TYR_CSK, the C-terminal phosphorylation motif in Src-family proteins targeted by the non-receptor tyrosine kinase Csk family^24^. Across 12 proteins with a known MOD_TYR_CSK motif in ELM, f13268 predicts residues in the motif with an AUROC of 0.97. Notably, the highest activation of f13268 occurs at the phosphorylated tyrosine residue (**Figure 3B**), indicating that this feature captures the biological characteristics of the motif. Importantly, MotifAE performs consistently across structural contexts, with median AUROCs of 0.88 in ordered regions and 0.84 in disordered regions, compared to 0.79 and 0.76, respectively, for SAE (**Figure 3C**; see Methods).

**Figure 3.**
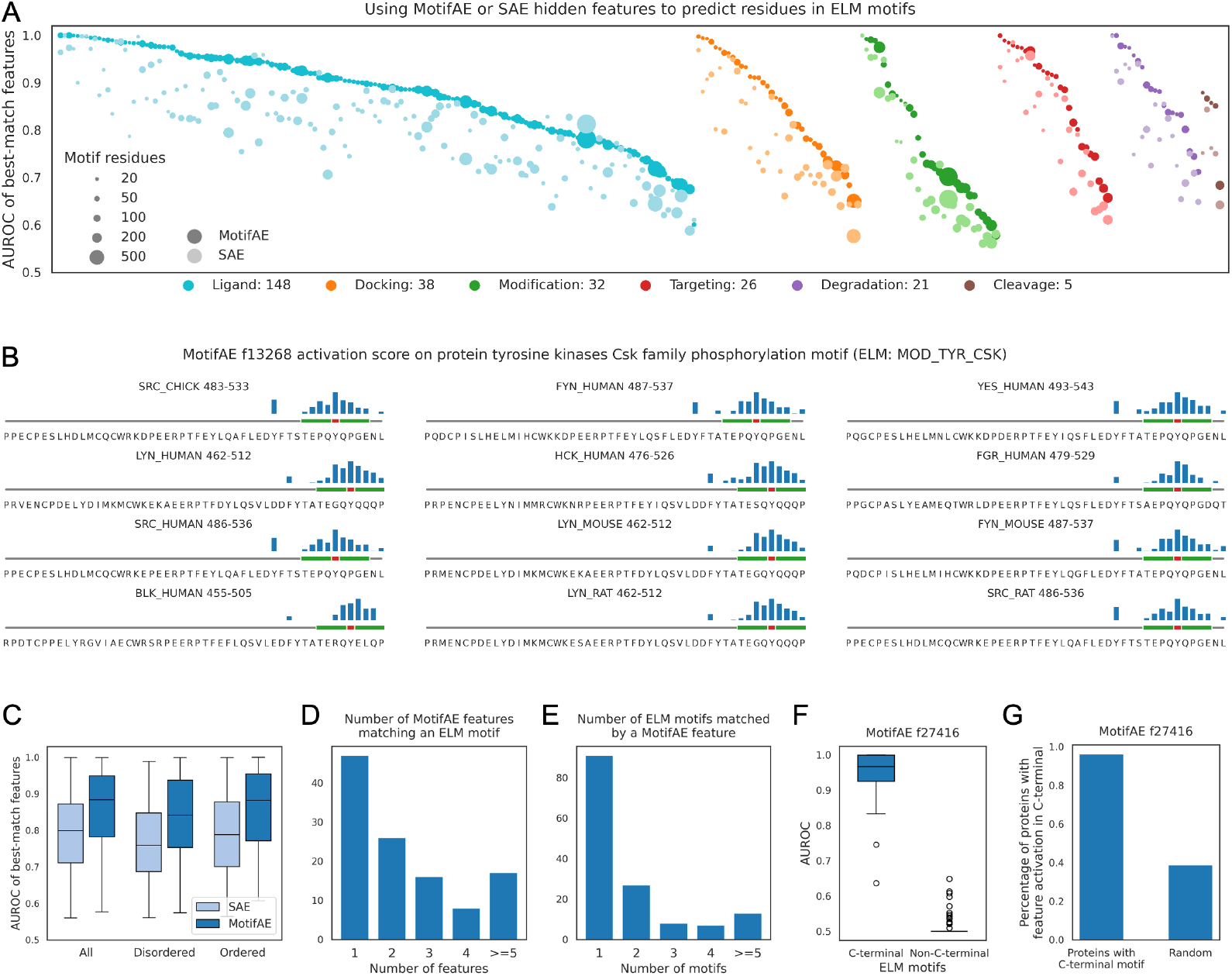
MotifAE captures known functional motifs. (**A**) AUROC of the best-match feature for each ELM motif. Each column shows the performance of MotifAE (dark dots) and SAE (light dots) for an ELM motif. Dot size indicates the number of residues in verified motif regions, and colors represent different ELM categories. The six categories and the number of motifs in each are listed below the plot. (**B**) Feature activation along protein sequences. The height of the blue bars indicates the normalized feature activation scores, the green region denotes the motif, and the red region marks the phosphorylated tyrosine residue. Only the C-terminal 50 residues of these proteins are shown for clarity. (**C**) Distribution of AUROCs of best-match features for ELM motifs. Disordered and ordered regions were classified using IUPred325. Only motifs with at least 20 residues located in disordered or ordered verified regions were included. The box extends from the first quartile to the third quartile, with a line marking the median, whiskers span to the most extreme points within 1.5× the interquartile range. (**D**) Number of features matching an ELM motif, (**E**) Number of motifs matched by a MotifAE feature, a motif– feature match is defined by AUROC > 0.9. (**F**) AUROC of MotifAE feature f27416 on C-terminal versus non-C-terminal ELM motifs. Non-C-terminal ELM motifs are defined as those with no verified regions within 10 residues of the C-terminus. (**G**) Percentage of proteins with activation of f27416 at the C-terminus. Proteins with known C-terminal motifs were obtained from the ELM dataset, random proteins were sampled from the evaluation set.

Subsequently, we analyzed the specificity of the relationship between ELM motifs and MotifAE features. We defined a matched motif–feature pair as one with AUROC > 0.9, resulting in 322 matched pairs involving 114 ELM motifs and 146 MotifAE features (**Figure S2**). Among these, 47 ELM motifs are matched by a single MotifAE feature, whereas 67 motifs are matched by multiple features (**Figure 3D**). On the feature side, 91 MotifAE features match only one motif (**Figure 3E**), whereas 55 features match multiple motifs. Notably, among the features that match multiple motifs, f27416 is one of the most prominent: it matches 17 motifs, all located at the C-terminus of proteins. Across all ELM motifs, f27416 achieves a median AUROC of 0.97 for 25 motifs exclusively located at the C-terminus but shows no predictive power for other motifs (**Figure 3F**). In addition, f27416 is not universally activated at the C-terminus of all proteins: among 320 proteins with known C-terminal motifs in ELM, 96% exhibit f27416 activation, compared with only 39% of randomly sampled proteins (**Figure 3G**).

Overall, these results demonstrate that the smoothness loss markedly improves MotifAE’s ability to capture functional motifs. These results also showed that MotifAE features exhibit varying levels of granularity: some correspond to specific functions, while others capture more general or widely shared functional patterns.

### The universe of motif sequence patterns captured by MotifAE

To characterize the sequence patterns of short motifs captured by MotifAE, we recorded activated regions in the representative protein dataset^22^ for each feature (see Methods), considering only regions with a length of 5–30 residues. Among the 40,960 features, 6,421 displayed coherent activations. Since functional motifs often exhibit distinct sequence preferences, we analyzed these activated regions and found that they exhibited significantly biased amino acid composition compared to random peptides (**Figure 4A**). Notably, approximately 400 features are almost exclusively activated by a single amino acid (**Figure 4B**; with one amino acid type accounting for >99% of all activations). To describe the sequence patterns, we constructed position-specific scoring matrices (PSSMs) for each MotifAE feature: activated regions (5–30 residues) were aligned using GibbsCluster^26^; the alignment cores, set to five residues for all features, were used to calculate the PSSMs (see Methods). The resulting PSSMs exhibited median KL divergence values (summed across the five residues) relative to the background of 10.7 (**Figure S3**). Among the PSSMs with well-defined sequence patterns, as indicated by higher KL divergence, some correspond to homopolymeric repeats (e.g., f13432 and f28775), while some exhibit multiple amino acid types (e.g., f2302, f36881, and f38747) (**Figure 4C**).

**Figure 4.**
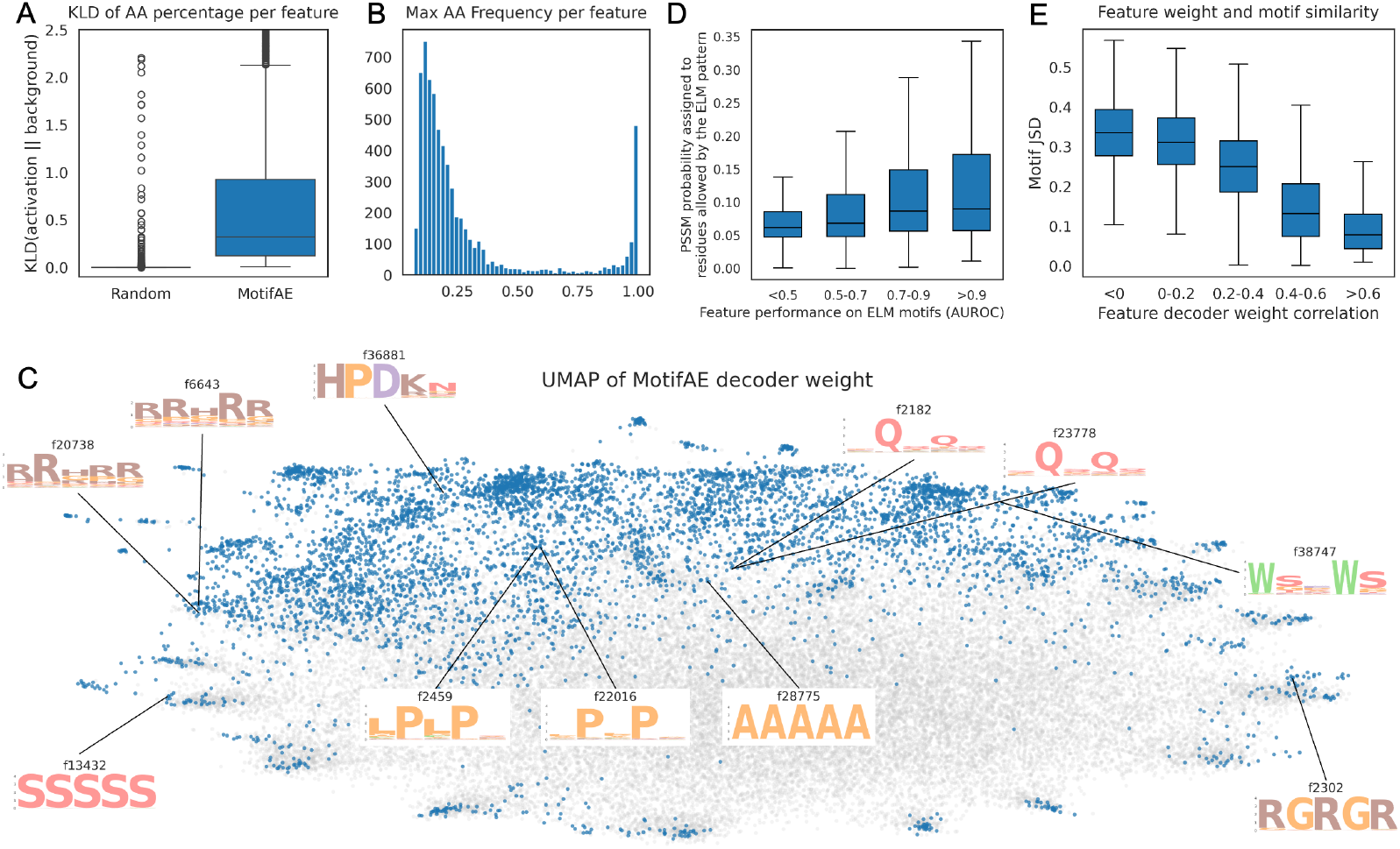
The universe of motif sequence patterns captured by MotifAE. (**A**) KL divergence between the amino acid composition of activated regions of each feature and the background (amino acid composition in the representative protein dataset). Random peptides matched in number and length distribution were used as the control. (**B**) Frequency of the most abundant amino acid in the activated regions of each feature. (**C**) UMAP visualization of MotifAE decoder weights. Each point represents a feature; gray points correspond to features without coherent activations, while blue points correspond to features with coherent activations. PSSM sequence logos are shown for some features. The x-axis shows the AUROC of using the feature activation score to identify ELM motif regions, and the y-axis shows the PSSM probability of residues permitted by the corresponding ELM regular expression. A value of 0.05 represents random expectation; higher values indicate greater similarity between the feature PSSM and the ELM regular expression (see Methods). (**E**) Relationship between feature similarity of decoder weights and similarity of PSSMs. Decoder weight similarity was measured using Pearson correlation, while PSSM similarity was measured using Jensen–Shannon divergence (JSD), where smaller JSD values indicate higher similarity.

To evaluate the validity of these PSSMs, we compared them against ELM motifs. Specifically, we aligned each feature PSSM to each ELM regular expression and analyzed the PSSM probabilities of amino acids permitted by the ELM expression (see Methods). We observed that MotifAE features that better identify motif regions also better match the corresponding ELM regular expressions (**Figure 4D**). Next, we analyzed the latent representation of each feature. In the MotifAE decoder, each feature has a weight vector that is multiplied by its activation score to reconstruct the embedding, thus, the decoder weights encode the properties of features in the embedding space. We found that features with similar decoder weight vectors tend to produce similar PSSMs (**Figure 4E**). For example, features f6643 and f20738 capture an “RRHRR” pattern; features f2459 and f22016 capture an “LPLP” pattern; and features f2182 and f23778 exhibit paired “Q” residues separated by a gap (**Figure 4C**). Together, by systematically characterizing the motif sequence patterns captured by MotifAE, we found that they exhibit specific sequence preferences, these sequence patterns align with known functional motifs and are reflected in the MotifAE weight space.

### MotifAE captures the functional organization of domains

Next, to investigate whether MotifAE captures longer sequence patterns, we assessed the ability of its feature activation scores to distinguish residues located inside versus outside annotated structural domains. We evaluated 400 CATH domains, including all major CATH classes: mainly α-helical, mainly β-sheet, mixed α/β, and others (see Methods). Considering the best-matching feature for each domain, MotifAE significantly outperforms the SAE across all CATH classes (**Figure 5A**). Additionally, we analyzed the specificity of the relationship between CATH domains and MotifAE features. We defined a matched domain–feature pair as one with AUROC > 0.8, resulting in 181 matched pairs involving 105 domains and 106 features. Among these, 71 domains are matched by a single MotifAE feature (**Figure 5B**). On the feature side, 84 MotifAE features match only one domain (**Figure 5C**). We note that this analysis is not intended to position MotifAE as a domain-identification method, as structure-alignment approaches already address that task effectively. Instead, this result highlight MotifAE’s ability to capture long-range sequence patterns.

**Figure 5.**
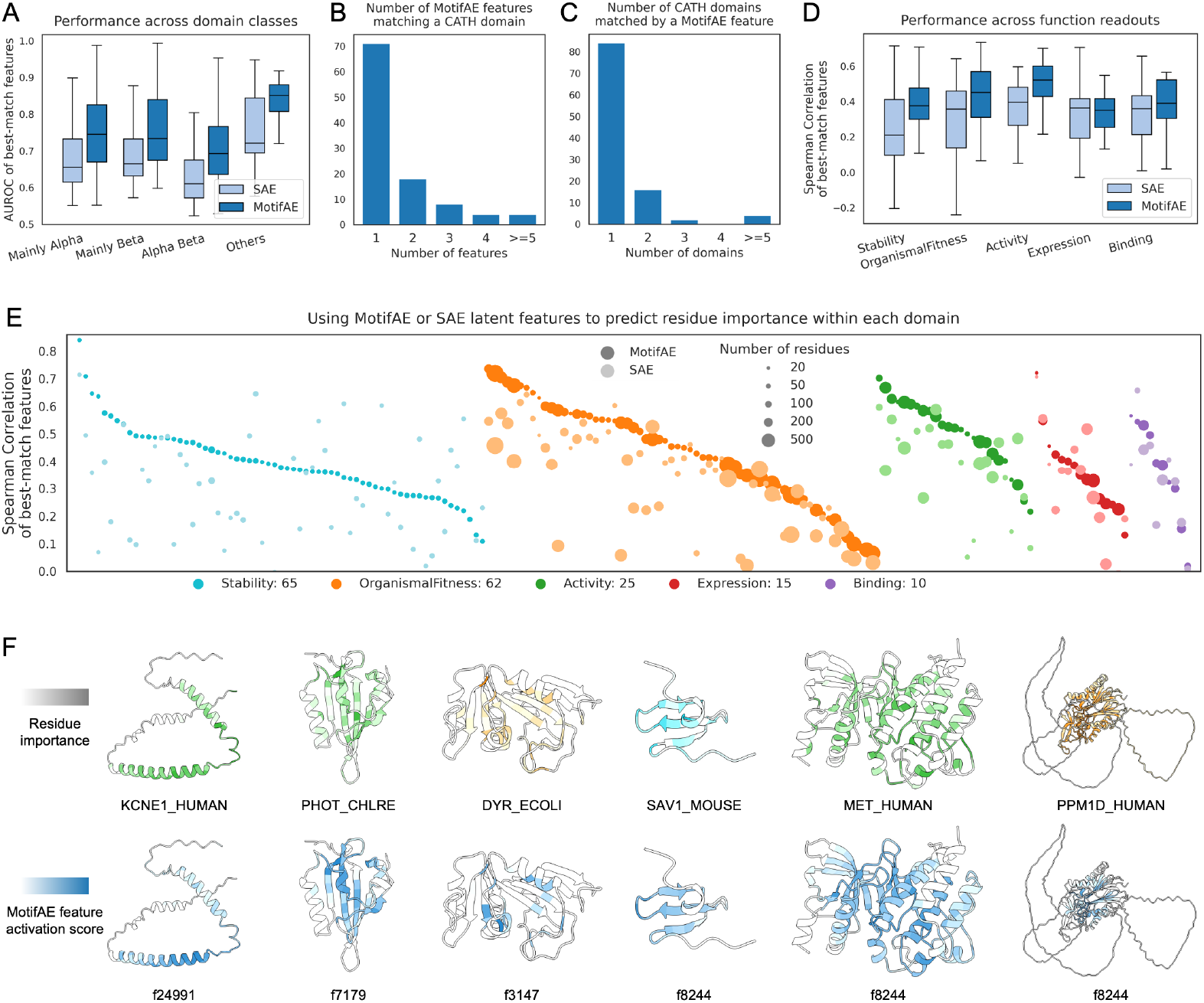
MotifAE captures the functional organization of domains. (**A**) Distribution of AUROCs for the best-match feature corresponding to each CATH domain, stratified by domain class. (**B**) Number of MotifAE features matching each CATH domain, and (**C**) Number of CATH domains matched by each MotifAE feature; a domain–feature match is defined as AUROC > 0.8. (**D**) Performance of the best-match features for each DMS experiment, stratified by ProteinGYM experimental categories. Performance is quantified using the Spearman correlation between the feature activation score and the mean mutation effect per residue, which serves as a proxy for residue importance across different functions. (**E**) Spearman correlations of the best-match feature for residue importance in each DMS experiment. Each column shows the performance of MotifAE (dark dots) and SAE (light dots) for a single experiment. Dot size reflects the number of residues with at least ten assayed mutations, and colors indicate experimental categories. The five categories and the number of experiments per category are listed below the plot. (**F**) Visualization of residue importance and MotifAE feature activations on protein structures. Upper panel: colors correspond to experimental categories, with darker colors indicating important residues with larger average mutation effects. Lower panel: darker colors denote higher feature activation scores.

Furthermore, we investigated whether MotifAE captures the functional organization of domains: the residue importance for different domain function. The mean mutation effect per residue measured by deep mutational scanning (DMS) was used as a proxy of residue importance for different functions. We analyzed 177 DMS datasets from ProteinGYM (see Methods), covering measurements of stability, organismal fitness (growth), activity, expression, and binding. Spearman correlation was used to quantify whether latent feature activation scores capture residue importance. Considering the best-match feature for each experiment, MotifAE achieves a median Spearman correlation of 0.41, significantly outperforming the standard SAE (0.33), with higher correlations observed for 80% of the experiments (**Figure 5D–E**). The improvement is particularly pronounced for residue importance for stability (median Spearman correlation 0.38 vs. 0.21), activity (0.52 vs. 0.40), and organismal fitness (0.45 vs. 0.36) (**Figure 5D**). Among individual features, f8824 is the most prominent, achieving a median Spearman correlation of 0.36 (**Figure S4**), slightly exceeding that of the best-match SAE features for each experiment. Examples of MotifAE features capturing the functional organization of domains are visualized on protein structures (**Figure 5F**, three examples are f8824), showing strong alignment between MotifAE latent feature activation scores and residue importance for different functions.

### Predicting a stability-specific fitness landscape via supervised MotifAE feature selection

pLM-predicted amino acid probabilities capture the protein fitness landscape and are widely used for mutation effect prediction and protein engineering. MotifAE retains essential information about fitness landscape (**Figure 2C**), and its amino acid probabilities are computed from reconstructed embeddings that integrate all latent features weighted by their activation scores (**Figure 1**), with different features capturing different functions and properties. Both pLM and MotifAE therefore predict a general fitness landscape that reflects combined constraints from multiple functional and biophysical requirements. Here, we investigated whether we can identify a subset of MotifAE features associated with a specific property, whether these features can be used to adjust predicted amino acid probabilities to reflect a property-specific fitness landscape, and whether this approach can improve prediction of that property and enable the rational engineering of domains with desired property levels.

We developed a supervised approach, called MotifAE-G, to identify features associated with a specific property from DMS experiments (**Figure 6A**). MotifAE-G introduces a binary gate to select which features contribute to the embedding reconstruction. The reconstructed embedding, based on the selected features, is then passed through the ESM2 MLP layer to predict amino acid probabilities and compute the LLR. A differentiable Spearman correlation between the LLR and the experimental measurement serves as the loss function. Importantly, this loss is used to update only the binary gate, while all parameters of ESM2 and MotifAE models remain fixed (see Methods). Following training, the binary gate identifies MotifAE features associated with the property of interest, and the residues activated by these features likely underlie the corresponding property.

**Figure 6.**
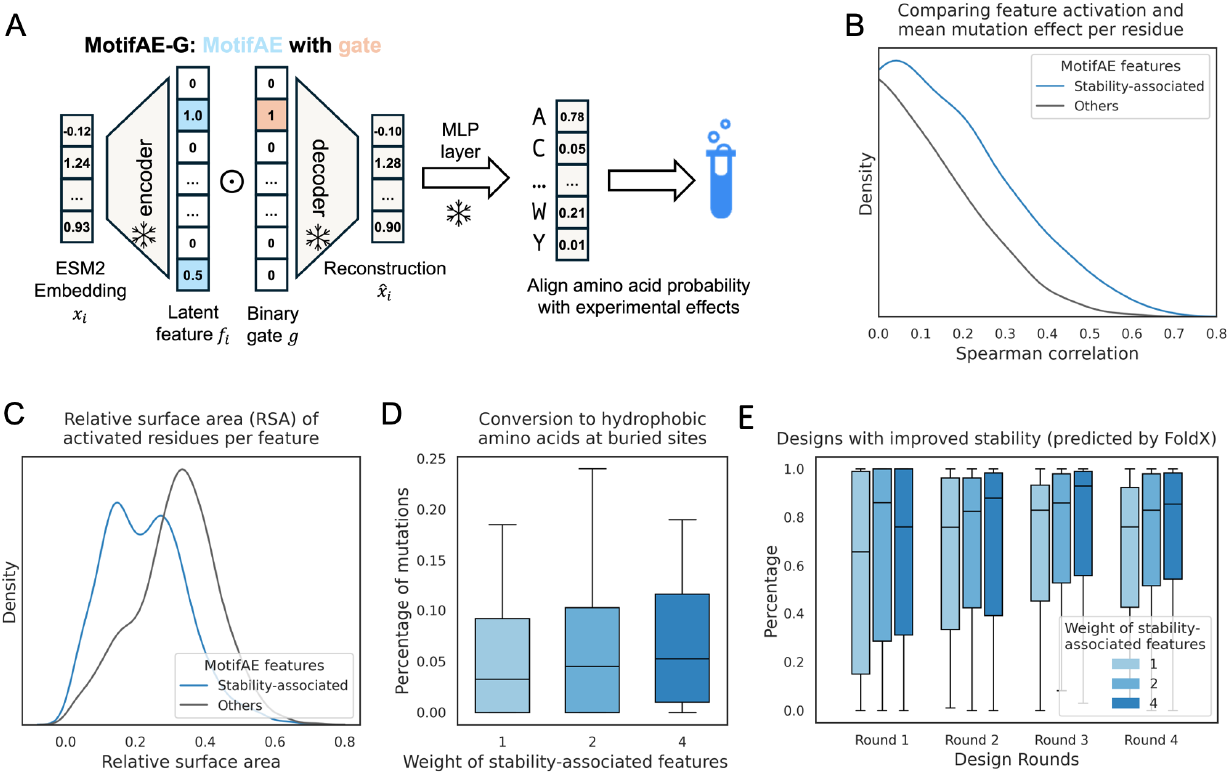
Predicting a stability-specific fitness landscape via supervised MotifAE feature selection. (**A**) Schematic of applying MotifAE-G framework to DMS data. A learnable binary gate was used to select latent features to align predicted LLR with experimental mutation effect; during training, only the gate parameters were updated. (**B**) Distribution of Spearman correlations between feature activation scores and mean mutation effects per residue across proteins. All features were compared against all DMS experiments, and only features activated in at least 30% of residues within a protein were included. Only positive correlations are shown. (**C**) Mean relative solvent accessibility (RSA) of activated residues for stability-associated versus other features, only features with at least 50 activated residues in 412 proteins were analyzed. (**D-E**) Designing proteins with improved stability using iterative redesign sampling. Gate weights of 1, 2, and 4 were applied to stability-associated features (shown in progressively darker blue), while other features were fixed at a gate weight of 1. (**D**) Buried residues were defined as sites with RSA < 0.2. All mutations from four rounds of redesign were included in the analysis. (**E**) Percentage of designs showing improved stability compared to their primary (unmutated) domains.

Specifically, we used a large-scale domain folding stability dataset^27^, focusing on DMS experiments of 412 domains (see Methods), including both natural and designed domains. Both ESM2 and MotifAE perform well on this dataset, with median Spearman correlations of 0.43 and 0.44, respectively (**Figure S5A**). To identify stability-associated features, we split the 412 domains into a 70% training set and a 30% test set, ensuring less than 30% sequence identity between them (see Methods). Using the MotifAE-G framework, we selected 1,404 stability-associated features from the training data. With these features, MotifAE-G achieves median Spearman correlations of 0.61 for both the training and test sets (**Figure S5B**). The improvement over ESM2 is more pronounced for domains lacking homologs in UniRef50^29^ (**Figure S5C**), the dataset used for ESM2 pretraining. To assess the biophysical relevance of selected features, we first evaluated how they capture residue importance for stability, finding that their activation scores correlated more strongly with mean mutation effect on stability per residue than those of non-selected features (**Figure 6B**). Next, we analyzed the relative solvent-accessible surface area and found that selected features preferentially activate at more buried residues (**Figure 6C**), which is consistent with the critical role of the domain core in maintaining structural stability.

Furthermore, we investigated whether steering selected features could enable the engineering of domains with enhanced stability. We employed an iterative redesign sampling strategy^30^: at each iteration, a single mutation was sampled according to amino acid probabilities predicted by MotifAE-G, with higher gate weights amplifying the influence of selected features, while other features kept a weight of one. We applied weights of 1, 2, and 4 to selected features, where a weight of 1 served as the baseline, equivalent to design using ESM2. Four rounds of mutations were applied to progressively refine the sequence, and the entire procedure was repeated 100 times for each domain (see Methods). We applied this strategy to test domains in the stability dataset. By analyzing the introduced mutations, we found that increasing the weights of stability-associated features lead to more conversions from other amino acids to hydrophobic amino acids at buried sites (**Figure 6D**), consistent with the fact that hydrophobic packing in the domain core is a key determinant of stability. We then evaluated the designed domains using FoldX^31^, a physics-based model that can estimate folding free energy (ΔG), applied to predicted structures^8^ (see Methods). We found that higher weights on stability-associated features yield more designs with improved stability relative to the original domains and larger improvements in predicted ΔG across the four design rounds (**Figure 6E, S5D**).

Overall, these results demonstrate that aligning latent features with experimental data enables MotifAE-G to identify meaningful, biophysically relevant features associated with domain folding stability. The selected MotifAE features support the prediction of a stability-specific fitness landscape that more accurately captures the effects of mutations on stability, generalizes across diverse proteins, and has the potential to facilitate the rational design of proteins with improved stability.

## Discussion

In this study, we present MotifAE, an unsupervised framework for extracting interpretable functional sequence patterns from pLMs. By incorporating a smoothness loss, MotifAE encourages coherent feature activation, which enhances its ability to discover functional motifs. This smoothness loss also propagates along the protein sequence, enabling the capture of longer sequence patterns, such as structural domains, and MotifAE feature activation scores correlate with residue importance for different domain functions and properties. Furthermore, we introduced MotifAE-G, a supervised extension that aligns latent features with experimental data, enabling the identification of features associated with a specific function or property. Beyond interpretation, this feature selection allows MotifAE-G to enhance predictive performance on related tasks and provides a strategy for the rational engineering of proteins.

While some SAE models have been trained for pLMs^16,18,32^, they often directly adopted strategies developed for large language models (LLMs). Our work introduces methodological advances in both training and interpretation. For training, we introduced an additional smoothness loss, defined as the L1 norm of latent feature differences between neighboring residues, sharing the same form as the sparsity L1 norm. Adding the smoothness loss preserves latent space sparsity, maintains reconstruction quality, and promotes coherent activation of latent features, facilitating identification of functional sequence patterns. For interpretation. A common approach is to compare activated residues with known functional sites, but this is limited by incomplete annotation databases and cannot reveal novel motifs. Although Simon et al.^32^ used LLMs to annotate latent features, their approach still relied on known protein functions. We provide two strategies for annotating and potentially discovering novel motifs. First, we calculate PSSM for each MotifAE feature; features with well-defined PSSM sequence patterns may correspond to novel motifs. Future studies are needed to systematically link these sequence patterns to biological functions and properties. Second, our MotifAE-G framework can be applied to annotate latent features using experimental datasets.

We applied MotifAE-G to the stability DMS dataset^27^ because it was generated in a single laboratory and is large enough to enable reliable identification of features associated with folding stability. In contrast, other DMS datasets usually target individual protein/domain and measure different properties, making them less suitable for feature selection. We note that the MotifAE-G framework can be applied more broadly. A general MotifAE-G application consists of two steps. First, a task-specific model is trained to predict experimental measurements from embeddings, with MotifAE reconstructions serve as a drop-in replacement for pLM embeddings; many existing models can be directly used (here, we used the ESM2 MLP layer). Second, a gate, which can be binary or continuous, selectively amplifies or attenuates MotifAE feature activations. This gate is trained on the same task to modulate both the reconstructed embeddings and the downstream predictions. Following training, the gate weights reflect the feature association with the measured function or property, and the activated residues of associated features may underlie the experimental measurement. Applying MotifAE-G to other large-scale experiments offers the potential to identify features associated with other biological functions and properties.

For benchmarking, we primarily compared MotifAE against the standard SAE. We note that Simon et al.^32^ already demonstrated that an SAE trained on pLM embeddings substantially outperforms both the raw pLM embeddings and an SAE trained on embeddings of pLM with shuffled weights in terms of interpretability. Therefore, we did not include those controls here, as MotifAE already outperforms the standard SAE. For motif region identification, no current model can perform systematic, unsupervised discovery of motifs, so no alternative methods are available for comparison. For residue importance for domain function and property, conservation scores derived from either multiple sequence alignments or pLM predictions perform comparably to, or even better than, best-match single MotifAE latent feature. However, these conservation scores are not decomposable: when a residue is predicted as conserved, the underlying reason—whether due to stability, binding, post-translational modification, or random noise—cannot be determined. In contrast, by selecting a subset of MotifAE features associated with specific properties, we can decompose the pLM-predicted conservation score into interpretable components, enabling better prediction of those properties.

Looking forward, MotifAE could be further improved in several directions. First, the current smoothness loss operates at the sequence level, incorporating three-dimensional proximity could enable the capture of more complex structural domains and their functional organization. Second, we currently use the L1 norm for the sparsity loss because it shares the same form as the smoothness loss. Future works are needed to explore how alternative sparsity-promoting strategies, such as TopK^33^ and BatchTopK^34^, can be integrated with the smoothness constraint. Third, since deep learning models inherently bias to their training data, both the pre-training dataset of the underlying pLM and the dataset used to train MotifAE influence the biological signals captured by latent features. Investigating how dataset composition shapes latent features is important. Fourth, MotifAE features exhibit different levels of granularity, understanding and controlling this granularity could improve feature interpretability and downstream utility. Lastly, our protein engineering results were evaluated only *in silico*; experimental validation will be important in the future to assess MotifAE’s potential for protein design.

Overall, we established a framework for the unsupervised discovery and interpretation of functional sequence patterns from pLMs, enabling the analysis of sequence patterns at evolutionary scale. Our work also contributes to the broader goal of making deep learning models in biology more interpretable. By systematically uncovering the sequence determinants of different protein functions and properties, MotifAE shows the potential to advance protein annotation, mutation effect interpretation, and rational protein engineering.

## Methods

### Sparse autoencoder architecture

Sparse autoencoder (SAE) projects language model embeddings into a sparse latent space. The model can be formulized as:

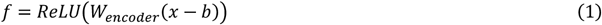

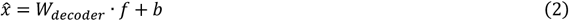

where *W*_*encoder*_ projects the original embedding *x* into the high dimensional latent space *f* and *W*_*decoder*_ does the reverse to get the reconstruction 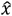. *b* is the bias term. ReLU was used as the activation function, setting negative values to zero and retaining positive values. Both the encoder and decoder consist of a single linear layer.

Here, the 650-million–parameter version of ESM2 was used, which produces embeddings with a dimensionality of 1,280. The SAE and MotifAE models were trained with varying latent space dimensions, and a dimension of 40,960 was selected (**Figure S1**). We trained MotifAE models using both normalized (along embedding dimension) and raw ESM2 embeddings and found that models trained on raw embeddings performed better, particularly for fitness prediction. Therefore, all main figures are presented using the MotifAE model trained on raw embeddings. In Figure S1, experiments using MotifAE models trained on normalized embeddings are presented.

### MotifAE loss function

MotifAE employs three losses: the reconstruction loss (*L*_*rec*_), the sparsity loss (*L*_*sp*_), and the smoothness loss (*L*_*smooth*_). The reconstruction and sparsity losses follow the SAE framework, preserving essential information from the input embeddings while enforcing sparse feature activations. The smoothness loss further encourages at least one nearby residue to have a similar latent feature activation, promoting coherent activations that capture the sequential continuity of protein motifs and basic structural elements. The loss functions are defined as follows:

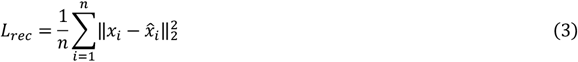

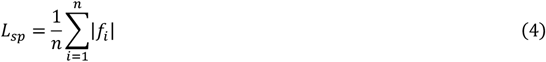

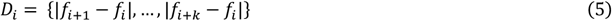

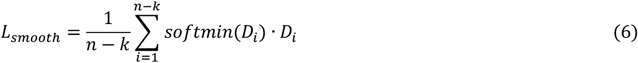

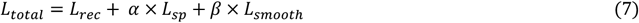

Here, *i* denotes the residue index; *n* denotes the protein length; *x* and 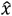 represent the raw and reconstructed embeddings, respectively; and *f* denotes the latent feature activations. *D*_*i*_ quantifies the L1-norm difference in latent feature activations between residue *i* and its *k* neighboring residues. *L*_*smooth*_ encourages at least one nearby residue to have a similar latent feature activation. To achieve this, we apply a SoftMin over *D*_*i*_ to approximate the minimum difference of latent feature activation. Only one direction along the sequence is considered for each residue, but since *L*_*smooth*_ is computed for the full sequence, the opposite direction is accounted for when evaluating neighboring residues. *k* is the local window size used for similarity calculation, which we set to three.

### Model training

MotifAE and SAE were trained and evaluated on 2.3 million representative protein sequences clustered from the AlphaFold Protein Structure Database using Foldseek^35^ (downloaded from https://afdb-cluster.steineggerlab.workers.dev/). Of these, 2.2 million sequences were used for training and the remainder for evaluation. Since ESM2 was trained with a maximum sequence length of 1,024, proteins exceeding this length were randomly truncated to a 1,024-residue region.

Models were implemented in PyTorch^36^ (v2.4) and optimized using the Adam optimizer with a maximum learning rate of 0.001 and a 500-step linear warm-up. A batch size of 40 proteins was used. Residue order was preserved during training to retain positional information required by the smoothness loss. MotifAE was trained with both sparsity and smoothness loss weights set to 0.4, whereas the standard SAE was trained with a sparsity loss weight of 0.85 and no smoothness loss. These loss weights were selected by testing multiple configurations to achieve a comparable number of activated latent features and similar reconstruction quality for both MotifAE and the standard SAE. The sparsity and smoothness loss weights were gradually annealed over 5,000 steps. Training was conducted for two epochs, and checkpoints at 80,000 steps for both MotifAE and SAE were used by considering latent sparsity and reconstruction quality. Each model required approximately 10 hours of training on a single NVIDIA L40S GPU.

### Mutation effect prediction and data processing

The wild-type marginal method^19^ was used to calculate log-likelihood ratio (LLR) to estimate mutation effects, which was calculated as:

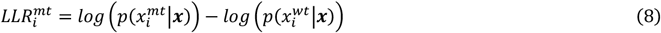

Where *i* represents residue index, *mt* and *wt* represent the mutant and wild-type amino acids, ***x*** is the full sequence without mask.

The ProteinGYM^20^ database comprises 216 deep mutational scanning (DMS) experiments spanning diverse functions and properties. In this study, only single substitution mutations were analyzed. For the domain functional organization analysis, mean mutation effects were computed for each residue, and only residues with at least ten assayed mutations were retained. Proteins containing fewer than twenty such residues were excluded, resulting in 180 DMS experiments. For each protein, only latent features that were activated in at least 30% of residues were analyzed. Three proteins lacking any SAE features meeting this threshold were excluded, yielding 177 proteins in the final analysis. The “DMS_score” values were inverted for Spearman correlation analysis, as smaller values in ProteinGYM correspond to larger mutation effects.

The ClinVar^23^ database provides expert-annotated mutations related to human diseases. Data preprocessing followed the procedure described in our previous work^37^. For this study, we considered only genes containing at least five pathogenic and five benign mutations, equal number of pathogenic and benign mutations were randomly sampled from each protein, resulting in a gene-balanced dataset of 2,272 pathogenic and 2,272 benign mutations in 207 genes. We used this dataset because ClinVar exhibits strong gene-level bias: the ratio of pathogenic to benign mutations varies substantially among genes^38^.

For protein stability^27^, the dataset “Tsuboyama2023_Dataset2_Dataset3_20230416.csv” and AlphaFold2-predicted protein structures were downloaded from https://zenodo.org/records/7992926. Proteins were defined using the “WT_name” column in the table. Only single substitution mutations with “ddG_ML” values were analyzed. The “ddG_ML” values for the same sequence were averaged. Homologs in the UniRef50 (downloaded in February 2025) were identified using MMseqs2^39^ search with the parameters: *-s 7 -a 1*. Solvent accessibility was computed from AlphaFold2-predicted structures using mdtraj^40^. The raw solvent-accessible surface areas were normalized by the maximum accessible area of each amino acid type, resulting in normalized values ranging from 0 to 1. For the domain functional organization analysis, the “ddG_ML” values were also inverted for Spearman correlation analysis (Figure 6B).

### Evaluation on ELM motifs and CATH domains

ELM motifs were downloaded from the ELM^1^ database in January 2025, and only instances annotated as *true positive* were analyzed. For proteins longer than 1,024 residues, a region of length 1,024 was selected with the motif region positioned at the center. IUPred3^25^ was used to predict disordered regions: residues with both long disordered region prediction > 0.5 and short disordered region prediction > 0.5 were classified as disordered, whereas all others were classified as ordered.

CATH^7^ v4.3.0 domain boundaries and annotations were downloaded. Sequences of PDB structure in CATH were mapped to UniProt^29^ Reviewed proteins, and the longest PDB structure was retained for each UniProt protein. Only UniProt proteins with lengths no longer than 1,024 residues were analyzed. CATH domains were required to lie fully within the mapped PDB–UniProt region, be at least 30 residues long, span no more than 70% of the full protein length, and be present in at least five different proteins. After filtering, 400 CATH domains across 5,076 proteins were included in our analysis.

Both tasks were formulated as a binary classification problem in which each residue was labeled as either belonging to a motif/domain or not. In MotifAE and SAE, each residue was represented by a 40,960-dimensional latent feature vector. For each latent feature, its activation value was used to predict motif/domain residues or not.

### Activated peptides and PSSM analyses

Activated peptides for each MotifAE feature were identified across the 2.3 million representative proteins. Only activated peptides with lengths between 5 and 30 residues were retained for motif-related analyses. To calculate position-specific scoring matrices (PSSMs), up to 1,000 activated peptides ranked by mean activation scores were used for each feature. GibbsCluster 2.0^26^, with parameters *-g 1 -l 5 -T -j 1 -I 1 -D 1*, was employed to align the activated peptides and remove outliers for each feature. The motif length was set to 5 amino acids. A trash cluster was used to remove sequences that did not align well with others. The aligned regions generated by GibbsCluster were subsequently used to construct PSSMs, which are calculated simply as the frequencies of 20 amino acids. PSSMs were visualized using the Logomaker^41^ Python package.

To compare ELM regular expression (regex) with PSSMs, each PSSM was slid along the regex to evaluate all possible alignments with at least three residues of overlap. For each alignment, the PSSM probabilities of residues permitted by the regex were analyzed, and the highest average across all valid alignments for each PSSM-regex pair was reported in Figure 4D.

### MotifAE-G architecture and training

The MotifAE-G introduced a gate to amplify or attenuate feature activations during embedding reconstruction, which were subsequently used for downstream tasks. The gate was applied to each latent feature of MotifAE as:

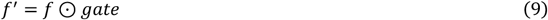

where ⊙ denotes elementwise multiplication.

For protein stability prediction, a learnable binary gate was introduced while keeping ESM2 and MotifAE parameters fixed. Each gate value could only take 0 or 1, and only features with a gate value of 1 were retained for reconstructing the embeddings. The gate was binarized using the Straight-Through Estimator^42^. In the forward pass, each gate was discretized as:

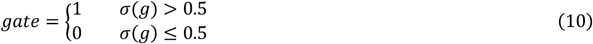

while in the backward pass, gradients were propagated directly through the sigmoid output σ(*g*).

The reconstructed embeddings from the selected features were passed through the ESM2 MLP layer to predict probabilities of 20 amino acids, which were then converted into LLRs between the mutant and wild-type amino acids (equation 8). The resulting LLRs were compared with mutation effect on protein stability “ddG_ML” to compute a soft Spearman correlation loss:

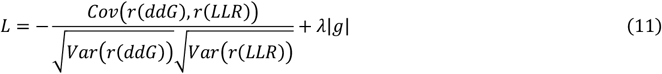

where *r*(·) denotes the differentiable rank function^43^. The first term maximizes the rank correlation between predicted and experimental mutation effects, while the second imposes an L1 regularization on the gate values to constrain the number of selected features. The weighting coefficient *λ* was set to 1 based on performance on evaluation set.

For MotifAE-G training, the 412 domains^27^ were clustered at 30% sequence identity, using MMseqs2^39^ easy-cluster with the parameters: *--min-seq-id 0*.*3 -c 0*.*5 --cov-mode 1*, yielding 189 clusters. A total of 285 domains from 133 clusters were used for training and validation, while 127 domains from the remaining 56 clusters were reserved for testing. 70% mutations in training proteins were used to select gates, while the remaining 30% were used as validation set to optimize hyperparameters.

Model training was implemented in PyTorch using the Adam optimizer with default parameters. Each batch contained a single protein, and gradients were accumulated across all training proteins before updating the gate parameters, meaning the parameters were updated once per epoch. Models were trained for 60 epochs, and the checkpoint with the highest validation Spearman correlation after 50 epochs was selected for downstream analyses.

### MotifAE-G protein design and stability prediction

pLMs can be used for protein design by sampling from their predicted amino acid probabilities. However, directly generating sequences from pLMs does not necessarily yield proteins with the desired property. To address this, several approaches have been proposed to fine-tune pLMs to guide the design process toward specific functions or properties^44,45^. Here, we explored whether MotifAE-G can be used for property-specific protein design. Specifically, we steered the stability-associated features identified from the DMS dataset to design proteins with improved stability.

Because ESM2 is not suitable for de novo sequence generation, we adopted an iterative redesign sampling strategy^30^. We performed four rounds of iterative design for each representative protein in 56 test clusters. We did not conduct more rounds because introducing a few new core interactions is typically sufficient to stabilize the protein, whereas excessive mutations may alter the native fold. In each round, the probabilities of all single substitutions were predicted using MotifAE-G, with different weights applied to stability-associated features. The wild-type sequence was used without masking, all mutations were ranked by their relative probability compared to the wild type: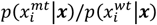. A top-k sampling strategy^30^ (with *k*=10 in this study) was applied to select one mutation, where the sampling probability of each mutation was proportional to its relative probability.

FoldX^31^ (version 20251231) was used to predict protein stability from the ESMFold^8^ predicted structures. Specifically, the “RepairPDB” command was first applied, followed by “Stability” to compute folding energy. The sign of the total energy was reversed to represent ΔG. Among the 56 representative proteins from the test clusters, 32 proteins with ESMFold pLDDT > 0.8 and FoldX-predicted ΔG > 2 kcal/mol were selected for subsequent design. Similarly, only designed sequences meeting the same thresholds were analyzed to ensure structural reliability and meaningful stability estimation.

## Data availability

All data used in this work are publicly available. 2.3 million representative proteins were downloaded from https://afdb-cluster.steineggerlab.workers.dev/. ProteinGYM DMS data were downloaded from https://proteingym.org/download. ELM motifs were downloaded from http://elm.eu.org/downloads.html. CATH domains were downloaded from https://www.cathdb.info/wiki?id=data:index. PDB-UniProt sequence matches were downloaded from https://ftp.ebi.ac.uk/pub/databases/msd/sifts/flatfiles/csv/uniprot_segments_observed.csv.gz. For protein stability, the DMS data and AlphaFold2-predicted protein structures were downloaded from https://zenodo.org/records/7992926.

## Code availability

Codes are available at GitHub: https://github.com/CHAOHOU-97/MotifAE.git. Model weights can be downloaded from: https://doi.org/10.5281/zenodo.17488191.

## Acknowledgments

This work was supported by NIH grants R35GM149527 and Simons Foundation SFARI #1019623.

## Contributions

Y.S. and C.H. conceived the study. C.H. designed and implemented the experiments. D.L. assisted with mutation analysis. All authors evaluated and interpreted the results and approved the final version of the manuscript.

## Supplementary information

**Figure S1.**
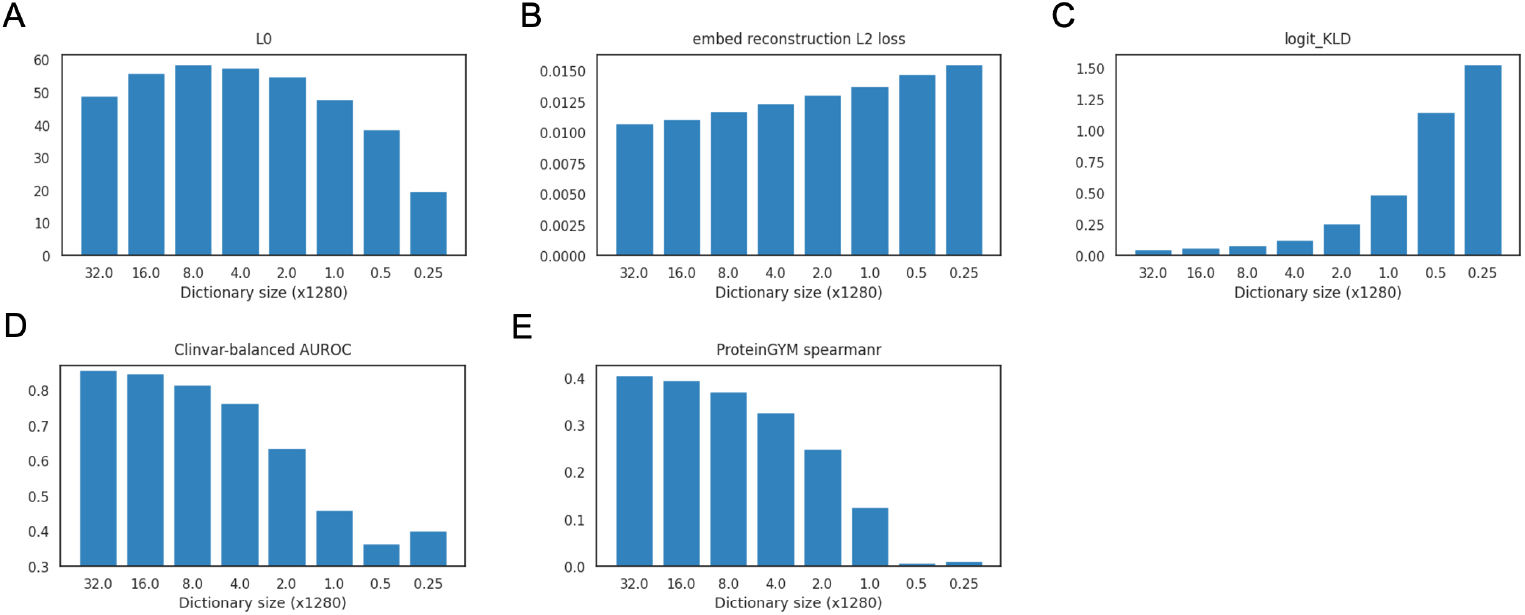
Comparison of different latent dimensions in MotifAE. The values on the x-axis represent the expansion ratio relative to the embedding dimension (1,280). We note that the results shown here are obtained from MotifAE trained on normalized ESM2 embeddings in our preliminary experiment, whereas the results in the main figures are MotifAE trained on embeddings without normalization. Therefore, the absolute values in some figures here may differ from the main figures, especially the reconstruction L2 loss, but the overall conclusions should remain consistent. (**A**) Average number of non-zero latent feature activations per residue on the evaluation set. (**B**) Reconstruction loss on the evaluation set. (**C**) Comparison of amino acid probability distributions predicted from MotifAE reconstructions with those predicted from ESM2 raw embeddings, both using the fixed ESM2 MLP prediction head. A KLD value of zero indicates identical distributions. (**D–E**) Performance on mutation prediction using the wild-type log-likelihood ratio. Mean Spearman correlation on ProteinGYM dataset was reported.

**Figure S2.**
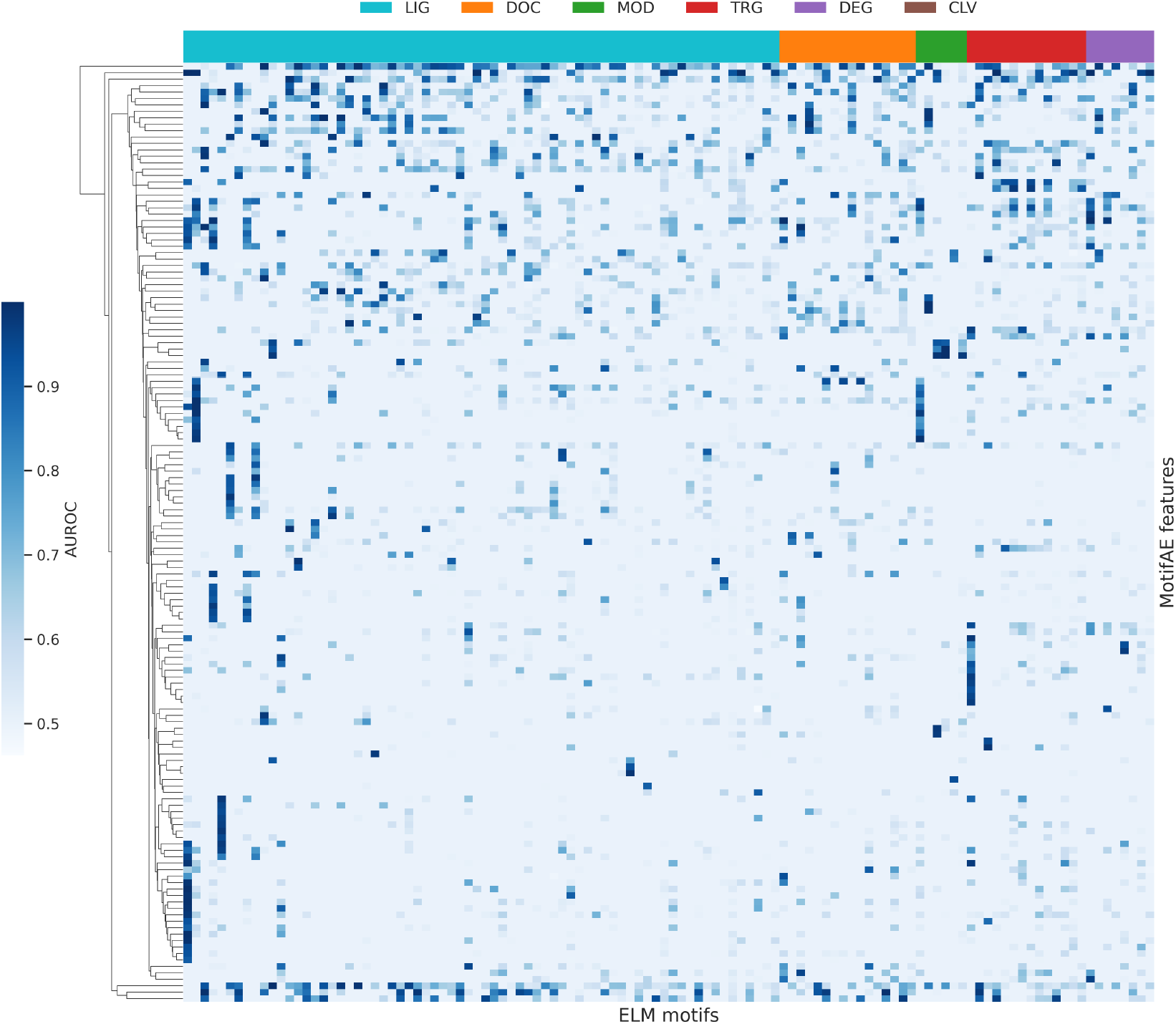
MotifAE feature performance on ELM motifs. Matched motif–feature pairs were defined as those with AUROC > 0.9, resulting in 322 matched pairs involving 114 ELM motifs and 146 MotifAE features. The heatmap shows the performance (AUROC, indicated by blue intensity) of these 146 features across all 114 motifs. The six motif categories on the x-axis are the same as those shown in Figure 3A.

**Figure S3.**
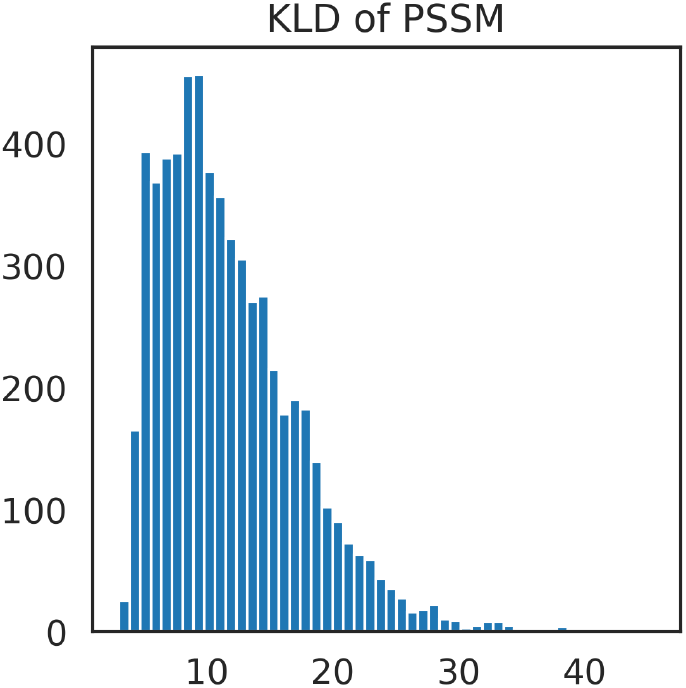
KL divergence of MotifAE feature PSSMs compared to the background amino acid frequency. KL divergence values were summed across the five aligned core positions.

**Figure S4.**
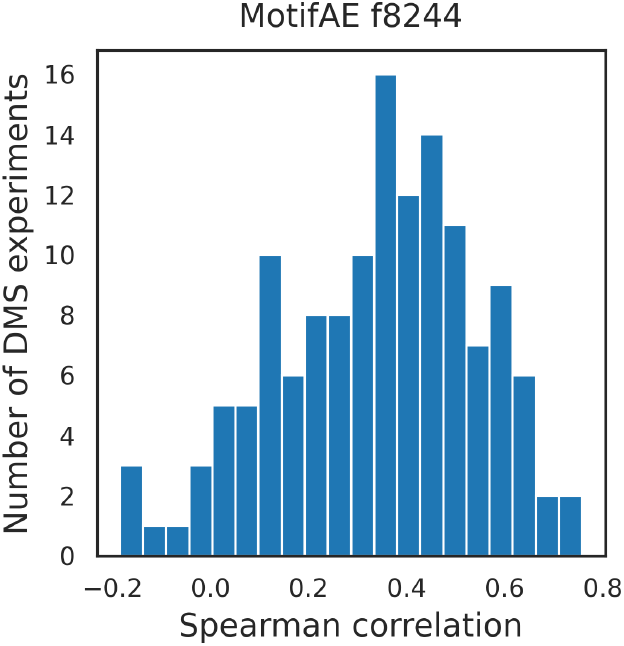
Spearman correlations of MotifAE feature f8824 in predicting the mean mutation effect per residue. A total of 139 DMS experiments with at least 30% of residues activated for f8824 were included in this analysis.

**Figure S5.**
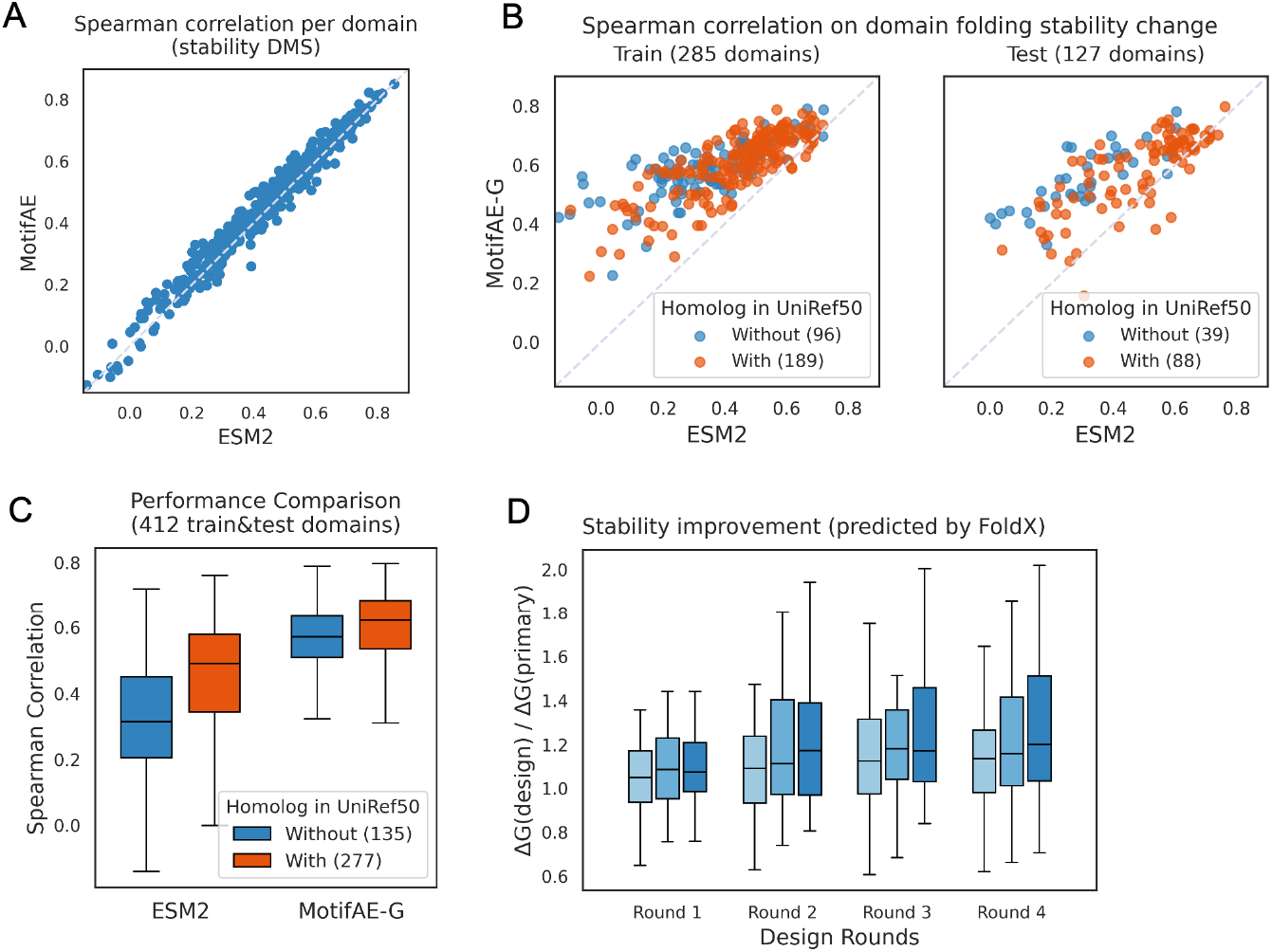
(**A**) Zero-shot performance (Spearman correlation) of ESM2 and MotifAE on 412 domain stability DMS datasets. (**B**) Comparison of MotifAE-G and ESM2 performance on training and test stability DMS datasets; each point represents a domain; color represents whether the domain has homolog in UniRef50 (ESM2 training set). (**C**) Comparison of MotifAE-G and ESM2 performance on domains with and without homolog in UniRef50. (**D**) Relative FoldX ΔG compared to the primary protein, 1 indicates no improvement. The median ΔG ratios across all designs for each protein in each round were shown.

## References

1. Kumar, M. et al. ELM-the Eukaryotic Linear Motif resource-2024 update. Nucleic Acids Res 52, D442–D455 (2024).

2. Tokheim, C. et al. Systematic characterization of mutations altering protein degradation in human cancers. Mol Cell 81, 1292–1308 e11 (2021).

3. Savojardo, C., Martelli, P. L. & Casadio, R. Finding functional motifs in protein sequences with deep learning and natural language models. Current Opinion in Structural Biology 81, 102641 (2023).

4. Hou, C., Li, Y., Wang, M., Wu, H. & Li, T. Systematic prediction of degrons and E3 ubiquitin ligase binding via deep learning. BMC Biol 20, 162 (2022).

5. Berman, H. M. et al. The Protein Data Bank. Nucleic acids research 28, 235–242 (2000).

6. Lau, A. M. et al. Exploring structural diversity across the protein universe with The Encyclopedia of Domains. Science https://doi.org/10.1126/science.adq4946 (2024) doi:10.1126/science.adq4946.

7. Waman, V. P. et al. CATH 2024: CATH-AlphaFlow Doubles the Number of Structures in CATH and Reveals Nearly 200 New Folds. Journal of Molecular Biology 436, 168551 (2024).

8. Lin, Z. et al. Evolutionary-scale prediction of atomic-level protein structure with a language model. Science 379, 1123–1130 (2023).

9. Jumper, J. et al. Highly accurate protein structure prediction with AlphaFold. Nature https://doi.org/10.1038/s41586-021-03819-2 (2021) doi:10.1038/s41586-021-03819-2.

10. Kulmanov, M. et al. Protein function prediction as approximate semantic entailment. Nature Machine Intelligence 6, 220–228 (2024).

11. Zhang, Z. et al. Protein language models learn evolutionary statistics of interacting sequence motifs. Proc Natl Acad Sci U S A 121, e2406285121 (2024).

12. Vig, J. et al. Bertology meets biology: Interpreting attention in protein language models. (2020).

13. Rives, A. et al. Biological structure and function emerge from scaling unsupervised learning to 250 million protein sequences. Proc Natl Acad Sci U S A 118, (2021).

14. Cunningham, H., Ewart, A., Riggs, L., Huben, R. & Sharkey, L. Sparse Autoencoders Find Highly Interpretable Features in Language Models. Preprint at 10.48550/arXiv.2309.08600 (2023).

15. Yun, Z., Chen, Y., Olshausen, B. A. & LeCun, Y. Transformer visualization via dictionary learning: contextualized embedding as a linear superposition of transformer factors. Preprint at 10.48550/arXiv.2103.15949 (2023).

16. Adams, E., Bai, L., Lee, M., Yu, Y. & AlQuraishi, M. From Mechanistic Interpretability to Mechanistic Biology: Training, Evaluating, and Interpreting Sparse Autoencoders on Protein Language Models. 2025.02.06.636901 Preprint at 10.1101/2025.02.06.636901 (2025).

17. Simon, E. & Zou, J. InterPLM: discovering interpretable features in protein language models via sparse autoencoders. Nat Methods 22, 2107–2117 (2025).

18. Gujral, O., Bafna, M., Alm, E. & Berger, B. Sparse autoencoders uncover biologically interpretable features in protein language model representations. Proceedings of the National Academy of Sciences 122, e2506316122 (2025).

19. Brandes, N., Goldman, G., Wang, C. H., Ye, C. J. & Ntranos, V. Genome-wide prediction of disease variant effects with a deep protein language model. Nat Genet 55, 1512–1522 (2023).

20. Notin, P. et al. ProteinGym: Large-Scale Benchmarks for Protein Fitness Prediction and Design. Advances in Neural Information Processing Systems 36, 64331–64379 (2023).

21. Hou, C., Liu, D., Zafar, A. & Shen, Y. Understanding Language Model Scaling on Protein Fitness Prediction. 2025.04.25.650688 Preprint at 10.1101/2025.04.25.650688 (2025).

22. Barrio-Hernandez, I. et al. Clustering predicted structures at the scale of the known protein universe. Nature 622, 637–645 (2023).

23. Landrum, M. J. et al. ClinVar: updates to support classifications of both germline and somatic variants. Nucleic Acids Res 53, D1313– D1321 (2025).

24. Okada, M. Regulation of the Src Family Kinases by Csk. Int J Biol Sci 8, 1385–1397 (2012).

25. Erdos, G., Pajkos, M. & Dosztanyi, Z. IUPred3: prediction of protein disorder enhanced with unambiguous experimental annotation and visualization of evolutionary conservation. Nucleic acids research 49, W297–W303 (2021).

26. Andreatta, M., Alvarez, B. & Nielsen, M. GibbsCluster: unsupervised clustering and alignment of peptide sequences. Nucleic Acids Res 45, W458–W463 (2017).

27. Tsuboyama, K. et al. Mega-scale experimental analysis of protein folding stability in biology and design. Nature 620, 434–444 (2023).

28. Hou, C., Zhao, H. & Shen, Y. Learning Biophysical Dynamics with Protein Language Models. bioRxiv 2024.10.11.617911 (2025) doi:10.1101/2024.10.11.617911.

29. UniProt, C. UniProt: a worldwide hub of protein knowledge. Nucleic Acids Res 47, D506–D515 (2019).

30. Darmawan, J. T., Gal, Y. & Notin, P. Sampling Protein Language Models for Functional Protein Design. in (2025).

31. Schymkowitz, J. et al. The FoldX web server: an online force field. Nucleic Acids Res 33, W382–8 (2005).

32. Simon, E. & Zou, J. InterPLM: discovering interpretable features in protein language models via sparse autoencoders. Nat Methods 22, 2107–2117 (2025).

33. Gao, L. et al. Scaling and evaluating sparse autoencoders. in (2024).

34. Bussmann, B., Leask, P. & Nanda, N. BatchTopK Sparse Autoencoders. Preprint at 10.48550/arXiv.2412.06410 (2024).

35. van Kempen, M. et al. Fast and accurate protein structure search with Foldseek. Nature Biotechnology http://dx.doi.org/10.1038/s41587-023-01773-0 (2023) doi:10.1038/s41587-023-01773-0.

36. Adam Paszke et al. PyTorch: an imperative style, high-performance deep learning library. in Proceedings of the 33rd International Conference on Neural Information Processing Systems Article 721 (Curran Associates Inc., 2019).

37. Zhong, G., Zhao, Y., Zhuang, D., Chung, W. K. & Shen, Y. PreMode predicts mode-of-action of missense variants by deep graph representation learning of protein sequence and structural context. Nat Commun 16, 7189 (2025).

38. Cheng, J. et al. Accurate proteome-wide missense variant effect prediction with AlphaMissense. Science https://doi.org/10.1126/science.adg7492 (2023) doi:10.1126/science.adg7492.

39. Steinegger, M. & Söding, J. MMseqs2 enables sensitive protein sequence searching for the analysis of massive data sets. Nat Biotechnol 35, 1026–1028 (2017).

40. McGibbon, R. T. et al. MDTraj: A Modern Open Library for the Analysis of Molecular Dynamics Trajectories. Biophys J 109, 1528–32 (2015).

41. Tareen, A. & Kinney, J. B. Logomaker: beautiful sequence logos in Python. Bioinformatics 36, 2272–2274 (2020).

42. Yin, P. et al. Understanding Straight-Through Estimator in Training Activation Quantized Neural Nets. Preprint at 10.48550/arXiv.1903.05662 (2019).

43. Blondel, M., Teboul, O., Berthet, Q. & Djolonga, J. Fast Differentiable Sorting and Ranking. Preprint at 10.48550/arXiv.2002.08871 (2020).

44. Dieckhaus, H., Brocidiacono, M., Randolph, N. Z. & Kuhlman, B. Transfer learning to leverage larger datasets for improved prediction of protein stability changes. Proceedings of the National Academy of Sciences 121, e2314853121 (2024).

45. Madani, A. et al. Large language models generate functional protein sequences across diverse families. Nat Biotechnol 41, 1099–1106 (2023).

